# MICAL2 Is a Super Enhancer Associated Gene that Promotes Pancreatic Cancer Growth and Metastasis

**DOI:** 10.1101/2024.06.26.600548

**Authors:** Bharti Garg, Sohini Khan, Deepa Sheikh Babu, Evangeline Mose, Kevin Gulay, Shweta Sharma, Divya Sood, Alexander T. Wenzel, Alexei Martsinkovskiy, Jay Patel, Dawn Jaquish, Guillem Lambies, Anthony D’Ippolito, Kathryn Austgen, Brian Johnston, David Orlando, Gung Ho Jang, Steven Gallinger, Elliot Goodfellow, Pnina Brodt, Cosimo Commisso, Pablo Tamayo, Jill P. Mesirov, Herve Tiriac, Andrew M. Lowy

**Affiliations:** Department of Surgery, Division of Surgical Oncology, Moores Cancer Center, University of California, San Diego. La Jolla, CA, USA; Department of Medicine, Moores Cancer Center, University of California, San Diego, La Jolla, CA, USA; Cancer Metabolism and Microenvironment Program, NCI-Designated Cancer Center, Sanford Burnham Prebys Medical Discovery Institute, La Jolla, CA, USA; Syros Pharmaceuticals, Cambridge, MA, USA; PanCuRx Translational Research Initiative, Ontario Institute for Cancer Research, Toronto, Ontario, Canada; Princess Margaret Cancer Centre, University Health Network, Toronto, Ontario, Canada; Department of Surgery, McGill University, Montreal, Quebec, Canada; Cancer Research Program, Research Institute of the McGill University Health Centre, Montreal, Quebec, Canada; Departments of Surgery, Oncology and Medicine, McGill University, Montreal, Quebec, Canada

## Abstract

Pancreatic ductal adenocarcinoma (PDAC) remains one of the deadliest solid cancers and thus identifying more effective therapies is a major unmet need. In this study we characterized the super enhancer (SE) landscape of human PDAC to identify novel, potentially targetable, drivers of the disease. Our analysis revealed that *MICAL2* is a super enhancer-associated gene in human PDAC. MICAL2 is a flavin monooxygenase that induces actin depolymerization and indirectly promotes SRF transcription by modulating the availability of serum response factor coactivators myocardin related transcription factors (MRTF-A and MRTF-B). We found that MICAL2 is overexpressed in PDAC and correlates with poor patient prognosis. Transcriptional analysis revealed that MICAL2 upregulates KRAS and EMT signaling pathways, contributing to tumor growth and metastasis. In loss and gain of function experiments in human and mouse PDAC cells, we observed that MICAL2 promotes both ERK1/2 and AKT activation. Consistent with its role in actin depolymerization and KRAS signaling, loss of MICAL2 expression also inhibited macropinocytosis. Through *in vitro* phenotypic analyses, we show that MICAL2, MRTF-A and MRTF-B influence PDAC cell proliferation, migration and promote cell cycle progression. Importantly, we demonstrate that MICAL2 is essential for *in vivo* tumor growth and metastasis. Interestingly, we find that MRTF-B, but not MRTF-A, phenocopies MICAL2-driven phenotypes *in vivo*. This study highlights the multiple ways in which MICAL2 impacts PDAC biology and suggests that its inhibition may impede PDAC progression. Our results provide a foundation for future investigations into the role of MICAL2 in PDAC and its potential as a target for therapeutic intervention.

## INTRODUCTION

Pancreatic ductal adenocarcinoma (PDAC) remains one of the most lethal malignancies worldwide, characterized by early metastasis, resistance to conventional therapies, and a dismal prognosis [1]. Despite notable progress in comprehending the genetics of PDAC, discerning transcriptional subtypes, and an increased understanding of its tumor microenvironment, efficacious treatment modalities remain elusive [2]. It is discouraging that our most effective therapeutic approaches still predominantly comprise traditional cytotoxic therapies, which often entail significant adverse effects on patients’ quality of life.

In recent years, extensive research has shed light on large, highly active chromatin regions known as "super-enhancers" (SEs), which play a critical role in defining cell identity and state in both normal and malignant cells [3, 4]. Previous studies have indicated that histone 3 lysine-27 acetylation (H3K27ac) marks serve as a reliable indicator for demarcating super-enhancers efficiently and robustly. These chromatin regions can regulate key genes that govern cell phenotype. In tumor cells, this regulatory mechanism may encompass both oncogene and non-oncogene drivers of the transformed state. We hypothesized that PDAC is driven and sustained by the activation of super-enhancers and that delineating the genes associated with these super-enhancers could unveil novel therapeutic targets for drug development. To investigate this hypothesis, we characterized the epigenetic landscape of normal and PDAC tissues obtained from primary patient samples. Using chromatin immunoprecipitation and sequencing (ChIPseq), we identified *MICAL2* as a super enhancer associated gene in human PDAC samples and confirmed its overexpression at the RNA and protein level in both human tissues, cell lines as well in murine models.

*MICAL2* is a member of the MICAL (molecules interacting with CasL) protein family, evolutionarily conserved flavin monooxygenases whose canonical function is the oxidation and resultant depolymerization of actin [5]. Unique to its other family members *MICAL1* and *3*, *MICAL2* has no autoinhibitory domain and is thus, constitutively active. MICAL2, which is present in both the cytoplasm and the nucleus, was previously shown to indirectly regulate serum response factor (SRF) mediated transcription through its modulation of nuclear G actin levels [6]. G actin acts to sequester myocardin related transcription factors A and B (MRTF-A and -B), coactivators of SRF. Nuclear accumulation of MRTFs is associated with the upregulation of genes associated with cell migration, fibrosis, and EMT, although MRTF-A has been the subject of most cancer-related studies [7]. Interestingly, the role of MRTF-B in oncogenesis is much more uncertain, with some studies linking it to mesenchymal and hepatocellular tumor progression, while a recent study concluded it acts as a tumor suppressor in human and murine colorectal cancer [8]. MICAL2 was first linked to malignant disease when its splice variants were found to be overexpressed in prostate cancer [9, 10]. More recently, studies have revealed that MICAL2 may promote EMT, migration, and invasion in the lung [11], gastric [12], and breast cancer [13], yet no prior studies have experimentally probed the role of MICAL2 in pancreatic cancer biology, nor comprehensively characterized MICAL2-regulated pathways and the specific roles of MRTF-A versus -B in oncogenic phenotypes. After we identified *MICAL2* as a highly ranked super-enhancer-associated gene in human PDAC, we found its expression correlated with poor prognosis in patients who had undergone surgical resection. We then determined that MICAL2 promotes PDAC growth and metastasis and that it is associated with patterns of gene expression associated with KRAS. We have identified a MICAL2/MRTF-B/SRF transcriptional program that underpins these malignant hallmarks. Our findings support the hypothesis that MICAL2 drives pancreatic cancer progression and that as an enzyme, it is worthy of further study as a potential therapeutic target for PDAC.

## MATERIAL AND METHODS

### Human Specimens

Normal and tumor pancreatic tissues were obtained from patients undergoing surgical resection or tissue biopsy at the University of California San Diego Health (UCSD Health). All tissue donations and experiments were reviewed and approved by the Institutional Review Board of UC San Diego Health. Written informed consent was obtained prior to acquisition of tissue from all patients. The studies were conducted in accordance with recognized ethical guidelines (Declaration of Helsinki). Samples were confirmed to be tumor or normal based on pathologist assessment (**Table S1**).

### Animals and *in vivo* procedures

All mouse protocols were reviewed and approved by the Institutional Animal Care and Use Committee of the University of California, San Diego. Five to eight weeks of age-matched and sex-balanced Male and Female F1 hybrid and nude mice were purchased from the Jackson Laboratory and used for each experiment. We used murine and human cell lines such as KPC46, AsPC-1, and BxPc-3 for in vivo experiments. KPC46 cells were injected in F1 hybrid mice, and AsPC-1 and BxPc3 cells were injected in NSG mice (JAX). For orthotopic and subcutaneous tumor challenges, mice were administered intra-pancreatic and flank injections of 50k KPC46 cells, 500k AsPC-1, and BxPc3 cells suspended in Matrigel. For the liver metastases model, we injected 50k KPC46 and 500k BxPC-3 cells in PBS via the intrasplenic route. Animals were euthanized after three weeks post-implantation for tumor weight and volume analysis.

### Cell Lines

KPC46 cell line was developed in our laboratory from *LSL-Kras^G12D/+^*;*LSL-Trp53^R172H/+^*;*Pdx-1-Cre* (KPC) mice, which develop spontaneous tumors [14]. AsPC-1 and BxPc3 cells were obtained from ATCC. All cell lines were maintained in Roswell Park Memorial Institute (RPMI) 1640 (Sigma-Aldrich, R8758) with 10% fetal bovine serum (FBS), penicillin/streptomycin at 37°C with 5% CO2, and all sh lines were maintained in Roswell Park Memorial Institute (RPMI) 1640 (Sigma-Aldrich, R8758) with 10% fetal bovine serum (FBS), penicillin/streptomycin and 5ug puromycin at 37°C with 5% CO2. We routinely checked for Mycoplasma contamination every two weeks. SiRNAs, Scramble control, and shRNAs against *MICAL2*, *MRTFA,* and *MRTFB* were obtained from Dharmacon^TM^ Horizon Discovery and Human and murine cell lines were infected using siRNA and shRNA using the protocol provided by the manufacturer. shRNA and siRNA sequences used in this study are summarized in **Tables S2 and S3.**

To establish BxPc3 overexpression (OE) and empty vector (EV) lines, the coding sequences of *MICAL2* (ENSG00000133816) and EGFP were cloned and inserted into the CSII-CMV-MCS- IRES2-Bsd vector, which was generously provided by the RIKEN BRC through the National BioResource Project of the MEXT, Japan (catalog number RDB04377) [15]. Following vector construction, the plasmid containing the lentiviral genome was transfected into HEK 293T cells along with helper plasmids, pCMV-VSV-G and pCAG-HIVgp, encoding viral envelope and packaging proteins. Viral supernatant was collected 2 days post-transfection and concentrated using Lenti-X lentivirus concentrator (Clontech). Cells were then infected with the concentrated lentivirus, and positive clones were selected by culturing cells in a medium containing 10 μg/ml blasticidin. The sequences of oligonucleotides used for CDS cloning can be found in **Table S4**. MICAL2 KD and OE efficiency was evaluated by using q-RTPCR.

### Quantitative RT-PCR

Total RNA from cell lines was extracted as per the manufacturer’s instructions (RNeasy Mini Kit, QIAGEN). Total RNA (1 μg) was reverse transcribed using the Quantitect Reverse Transcription Kit (Qiagen). Subsequently, specific transcripts were amplified by SYBR Green PCR Master Mix (USB) using a Bio-Rad CFX96 Real-Time System thermocycler. Where fold expression is specified, the comparative CT method was used to quantify gene expression. Expression was normalized 18S. Primers used for QPCR are detailed in **Table S5.**

### Western blotting

Western blotting was conducted following established protocols with slight adjustments. Initially, cell pellets were lysed in 2% SDS buffer supplemented with complete Protease (Roche, 11697498001) and PhosSTOP phosphatase (Roche, 4906845001) inhibitors after thorough washing with Dulbecco PBS. Subsequently, cell lysates were prepared using RIPA buffer, and protein concentrations were determined using the Pierce BCA Protein Assay Kit (Thermo Fisher Scientific, 23225). The proteins were denatured by adding NuPAGE LDS Sample Buffer (4×; Thermo Fisher Scientific, NP0008) and heating at 98°C for 10 minutes. For human and murine 2D cell lines, ten micrograms of proteins were separated by SDS-PAGE and transferred onto 0.2 μm PVDF membranes (Bio-Rad Laboratories, Inc., 1704156) using the Trans-Blot Turbo Transfer System (Bio-Rad Laboratories, Inc.). Following blocking with 5% skim milk or 5% bovine serum albumin in Tris-buffered saline with 5% TBST for 1 hour at room temperature, membranes were incubated overnight at 4°C with the primary antibody diluted in the blocking solution. After washing three times with TBST, membranes were incubated with the corresponding secondary anti-mouse (Thermo Fisher Scientific, 31430) or anti-rabbit IgG antibody (Thermo Fisher Scientific, 31460) in a blocking solution. Signals were visualized using Immobilon Western Chemiluminescent HRP substrate (MilliporeSigma, WBKLS0500) and captured using the ChemiDoc Imaging System (Bio-Rad Laboratories, Inc.). Data were analyzed using ImageJ and Image Lab software (17). The list of antibodies used for Western blotting is detailed in **Table S6.**

### Histology and immunohistochemistry

Tissues were fixed in 10% neutral-buffered formalin, embedded in paraffin, and sectioned at a thickness of 5 μm. Standard hematoxylin and eosin staining procedures were employed for histological examination. For immunohistochemistry (IHC), heat-induced antigen retrieval was achieved by immersing the tissue sections in citric acid buffer (pH 6.8) and subjecting them to pressure cooking for 20 minutes. Endogenous peroxidases were blocked using 3% H2O2 in tris-buffered saline (TBS) for 20 minutes at room temperature. To minimize nonspecific binding, tissue sections were incubated with a blocking solution containing 10% goat and 2.5% horse sera. Subsequently, the sections were exposed to the primary anti-MICAL2 antibody (13965-1-AP) overnight at 4°C. Following three washes with TBS containing 1% Tween 20 (TBST), the sections were treated with a Goat anti-Rabbit IgG ImmPRESS secondary antibody for 30 minutes at room temperature. After additional washing steps with TBST, the signals were developed using ImmPACT DAB HRP Substrate. Finally, the slides were counterstained, dehydrated, and coverslipped according to standard protocols.

### In vitro assays

Cell viability and proliferation rates were assessed using a CCK-8 proliferation kit. Initially, cells were seeded in 6-well plates at a density of 500k cells per well and transfected with specific siRNAs (siControl, siMical-2, siMRTF-A, siMRTF-B). After 72 hours, the cells were split and seeded in 96-well plates at a density of 1k cells per well for KPC46 and 5 × 103 cells per well for AsPC-1 in 5% FBS (Omega Scientific). Following incubation for 3, 4, 5, 6,7, and 8 days, 10 μl of CCK-8 reagent (Abcam, ab228554) was added to each well at each time point. Subsequently, the 96-well plate was incubated in the dark for 1 hour and then the absorbance was measured at 460 nm using a microplate reader (Biotek, Synergy HT). For the *in vitro* wound healing assay, human and mouse PDAC cell lines were cultured in 6-well plates until reaching 80–90% confluence. A straight-line scratch was created across the cell monolayer using a 1000 µL pipette tip. The cells were then rinsed three times with 1x PBS to eliminate unattached cells and debris before adding RPMI supplemented with 10% FBS. Following the acquisition of the initial image (0 hr), the plate was placed in a 37°C, 5% CO2 incubator. Subsequent images were captured at various time points (12, 16, 21, 32 and 72 hrs) using a phase contrast microscope. For the cell cycle assay, 1 million cells/well were seeded in a 6-well plate. The following day, cultures were washed with PBS and harvested by trypsinization. Cells were then washed again with ice-cold PBS to remove excess media and fixed with 6 ml of 70% ice-cold ethanol for 24 hours at −20°C. After fixation, tubes were centrifuged at 2000 RPM for 10 minutes, followed by washing with ice-cold PBS. The cells were suspended in 250 µl of TBS-T and treated with 50 µl of a 100 µg/ml stock of RNase and 200 µl of PI from a 50 µg/ml stock solution. Tubes were then incubated at 37°C in the dark for 15 minutes before analysis by FACS. For the SRF reporter assay, cells were transduced with Serum Response Factor (RhoA/ELK1/SRF) response element luciferase reporter lentiviral particles according to the manufacturer’s protocol (G and P Biosciences). Positive cells were selected using antibiotic Zeocin and Luciferase reporter assay was performed using Dual-Glo luciferase assay kits (Promega) according to the manufacturer’s instructions. Firefly luminescence was normalized by the viability and number of cells.

### Gelatin degradation assay and immunofluorescence

The gelatin degradation assay was performed essentially as described previously [16]. Briefly, 12-mm-diameter glass coverslips were presoaked in 70% ethanol for 20 minutes before use, transferred to a 24-well plate (Corning) and incubated at room temperature with 500μL poly-L-lysine (Sigma, P8920):PBS (Wisent) at a 1:1 ratio for 20 minutes. After incubation and washing thrice with 500μL PBS, 500μL of freshly prepared 0.5% glutaraldehyde in PBS was added to the wells and incubated for 20 minutes at room temperature to activate the poly-L-lysine surface for further protein binding. Coverslips were washed thrice with PBS and coated with a mixture of pre-warmed (37°C) 0.2% gelatin and FITC-gelatin (ThermoFisher, G13187) at a 6:1 ratio, washed and incubated with a fresh solution of 5 mg/ml sodium borohydride (Sigma, 452882) for 20 min to quench free aldehydes. After extensive washing, the coverslips were incubated in 24-well plates containing 70% ethanol and stored at 4 °C in the dark until used. The ethanol was removed, and coverslips were washed thrice with PBS before use. Coverslips were mounted with the ProLongTM gold antifade reagent (Thermo Fisher Scientific, Quebec, Canada) before image acquisition.

### Confocal microscopy and image analysis

Images were acquired using the Zeiss LSM780 laser scanning confocal microscope equipped with X63 oil objective. Five representative images were captured per cell type. Gelatin degradation appeared as punctate or diffuse ‘dark clearings’ (depending on incubation time) in the bright fluorescent gelatin due to loss of fluorescence signal. Gelatin matrix degradation was quantified by ImageJ. Data processing was performed using GraphPad Prism 6 and statistical analyses were performed using the one-way ANOVA.

### Macropinocytosis assay and quantification

Macropinocytosis assay and quantification were performed as previously described [17]. Briefly, cells transfected with a scrambled siRNA (siCTRL) or siRNA targeting MICAL2 (siMICAL2) were seeded on acid-washed glass coverslips in 24-well plates and subjected to serum and glutamine starvation for 24 hours. Media was then removed from the wells and the same media containing 1 mg/mL FITC-Dextran (ThermoFisher Scientific) was added back to the cells and plates were incubated for 30 minutes at 37°C. Thereafter, 4-5 washes with cold PBS were done and cells were fixed in 3.7% formaldehyde for 15 minutes. Nuclei were next stained with 2 μg/ml of DAPI (MilliPore) and coverslips were mounted on glass-slides with DAKO-fluorescent media (Agilent Technologies). Images were automatically captured at 40X magnification using a BioTek Cytation 5 (Agilent Technologies) and were subjected to automated analysis using the BioTek Gen5 software to calculate the relative macropinocytic index.

### RNA Sequencing

AsPC-1 Shcontrol and ShMICAL2 cells were cultured in 10-cm dishes in triplicates. Total RNA was extracted using TRIzol following the manufacturer’s instructions. RNA samples were submitted to Azenta Life Sciences for further analysis. Quality testing was carried out by measuring RNA integrity and OD readings (260/280 and 260/230) to have RIN scores of 9.4 or higher, and OD readings were within the 1.8 to 2.2 range. Differential expression analyses were analyzed by gene set enrichment analysis (GSEA) for Hallmarks.

### H3K27ac ChIP-Seq

Frozen tissue samples were homogenized using the Covaris CP02 cryoPREP Automated Dry Pulverizer. Homogenized tissue was fixed rotating at room temperature for eight minutes in 1% formaldehyde (ThermoFisher catalog #28906), diluted in PBS. Fixation was quenched with 1/20^th^ volume of 2.5M glycine. All buffers after fixation and prior to wash buffer before elution included HALT protease inhibitor (Life Technologies catalog #78438) and 1mM sodium butyrate to inhibit histone deacetylase activity.

Samples were pelleted and washed twice with cold PBS. Samples were then lysed rotating at 4°C with ChIP Lysis Buffer 1 (Boston BioProducts catalog # CHP-126) for ten minutes, then with Lysis Buffer 2 (Boston BioProducts catalog # CHP-127). Samples were then washed with Sonication Buffer (CHP-133). Samples were resuspended in sonication buffer and sonicated to an average size of 100-500 bp on the Covaris E220 focused ultrasonicator, in a volume of 750 µL sonication buffer. Following sonication, samples were collected and centrifuged at 20,000x*g* for 10 minutes at 4°C to pellet insoluble material. The supernatant was collected and diluted in one volume of ChIP Dilution Buffer (Boston BioProducts #CHP-143), and input controls were collected.

Chromatin was immunoprecipitation overnight (12-16 hours) rotating at 4°C with anti-H3K27ac antibody (Abcam, catalog #4729) conjugated to protein G Dynabeads (Life Technologies, catalog #10004D). The following day, immunoprecipitated chromatin was washed twice with low salt wash buffer (Boston BioProducts #CHP-146), once with high salt wash buffer (Boston BioProducts #CHP-147), once with LiCl wash buffer (Boston BioProducts #CHP-149), and once with Tris-based wash buffer (Boston BioProducts #CHP-148). Chromatin was eluted with ChIP Elution Buffer (Boston BioProducts #CHP-153) for one hour at 65°C with agitation at regular intervals. Eluted chromatin was reverse crosslinked for 16 hours at 65°C. Chromatin was then diluted in one volume of TE buffer, then treated with RNAse (Fisher catalog # AM2286) for 2 hours at 37°C and proteinase K (Fisher catalog # AM2548) for 1 hour at 55°C. An equal volume of phenol/chloroform (Fisher, catalog #15593031) was added to each sample, and the aqueous phase was collected in a Phase Lock Gel tube (VWR catalog #10847-802). Each sample received 1 mL ethanol, 20 µL 5M NaCl, and 30 µg GlycoBlue (Fisher Scientific, catalog # AM9516), which were then mixed by inversion and stored at −20°C overnight. The following day, the samples were centrifuged at 20,000x*g* for 20 minutes to precipitate the purified DNA. Pellets were washed with 800 µL cold 80% ethanol and a second centrifugation at 20,000x*g* for 10 minutes. Pellets were briefly air dried and resuspend in 50 µL water.

### H3K27ac Sequencing

Libraries were generated using the Swift Accel-NGS 2S Plus DNA library kit (Swift Biosciences catalog #21024). Samples were multiplexed six libraries per lane on an Illumina HiSeq 2500.

### Differential H3K27ac enrichment analysis

H3K27ac ChIP-seq reads were aligned to the hg19 human reference genome with bowtie2 (v2.0.5) using the option “--sensitive” [18]. Regions enriched for H3K27ac were called with MACS2 (v2.0.10) [19], using input-control data as background. To calculate differences in H3K27ac signal in PDAC samples compared to normal pancreas samples, we performed the following analyses steps: 1) Peaks with a MACS2 score < 20 were removed, and the remaining peaks were merged across samples to generate a universal peak set; 2) ChIP-seq reads and input-control reads overlapping each peak were tallied for each sample; 3) After applying a normalization factor generated using NCIS [20], input-control reads were subtracted from ChIP-seq reads for each peak and sample; 4) The resulting peaks were further filtered to remove those that did not have at least 20 normalized reads in at least two samples; 5) Batch effects were estimated and removed with a negative binomial generalized linear model via DESeq2; 6) Normalized read counts across technical replicates were averaged (mean) and rounded to the nearest integer; 7) These values were then used to calculate differences between PDAC and normal pancreas using DESeq2 [21].

### H3K27ac gene-set analyses

To determine if differential H3K27ac regions were enriched near genes belonging to specific biological processes or modes of regulation, we followed the framework described by McLean, et. al (GREAT [22]). Briefly, we performed the following analysis steps: 1) Regulatory domains were defined using GENCODE v19 protein-coding annotations with level 1 or 2 support; 2) Test peak sets were defined as peaks with an adjusted p-value ≤ 0.01 and a log2(fold-change) of either > 0 or < 0. The background peak set was defined as any region tested by DESeq2 as described above; 3) The width of each peak was adjusted to 1000 bp by extending the boundaries +/- 500 bp from the peak’s midpoint; 4) The enrichment of the test peak sets in a given gene set’s regulatory domains were calculating using “phyper” in R; 5) Multiple hypothesis testing correction was performed by the Benjamini-Hochberg method with “p.adjust”.

### Differential super-enhancer analysis

To call super-enhancers, we followed previously described methods [3, 23]. Briefly, we performed the following analysis steps: 1) MACS2 peaks were filtered using a p-value threshold of ≤ 1x10^-9^; 2) MACS2 peaks located within 12.5 kb of one another were merged to yield a set of “stitched” regions; 3) Stitched regions were merged across samples to generate a universal region set; 4) ChIP-seq reads and input-control reads overlapping each region were tallied for each sample; 5) After applying a normalization factor generated using NCIS [20], input-control reads were subtracted from ChIP-seq reads for each region and sample; 5) A negative binomial distribution was fit to these normalized region counts for each sample; 6) Regions were called as super-enhancers for a given sample if their counts were ≥ the 97.5^th^ percentile of the abovementioned estimated distribution. To determine which super-enhancers had differential H3K27ac enrichment in PDAC compared to normal pancreas, we performed the following analysis steps: 1) A universal set of super-enhancers was defined as the union of regions called as super-enhancers in four out of six normal pancreas samples, and five out of seven PDAC samples; 2) Using the quantifications for these super-enhancers, batch effects were estimated and removed with a negative binomial generalized linear model via DESeq2; 3) Normalized read counts across technical replicates were averaged (mean) and rounded to the nearest integer; 4) These values were then used to calculate differences between PDAC and normal pancreas using DESeq2 [21]. Library sizes were determined using the full stitched region set.

### H3K27ac enrichment visualization

To visualize input normalized H3K27ac signal across the genome, we generated bigwig files for each sample using “bamCompare” from deepTools2 [24]. More specifically, we used the options: “--operation subtract”, “--scaleFactors”, “--binSize 10”, “--smoothLength 30”, “—normalize “Using None”, and “--extendReads”. To determine scaling factors, we used NCIS [20] and further adjusted these factors by the number of mapped reads (in millions) of the ChIP-seq sample. We estimated the fragment size to extend the reads using “phantompeakqualtools” [25]. A signal was averaged (mean) for technical replicates of the same biological sample.

## Statistical analysis

GraphPad Prism version 9 was used for the graphical representation of data. Results are expressed as the mean and s.e.m. unless otherwise indicated in the figure legend. For the tumor growth studies, the data shown represent independent experiments with biological replicates. A Student’s two-tailed unpaired *t*-test was used for the experiments with two groups unless otherwise indicated. For multiple group comparisons, a one-way or two-way ANOVA was performed followed by a Tukey test unless otherwise indicated in the figure legend. Data were considered significant if *P* < 0.05; exact values are shown in each figure legend.

## Data availability

The analyzed data, H3K27ac ChIP-sequencing (ChIP-seq) and RNA sequencing (RNA-seq) generated in this study will be publicly available in the GEO database. All other data are available upon request from the corresponding authors.”

## RESULTS

### MICAL2 is a super enhancer-associated gene in human PDAC

We sought to identify tumor-specific active super-enhancer (SE) regions in human PDAC. We obtained tumor (n = 7) from surgical resections of PDAC and normal pancreatic tissues (n = 6) from patients undergoing resection for non-adenocarcinoma histology (**Table S1**). Tissues were lysed and we performed histone 3 lysine 27 acetylation (H3K27ac) ChIP-seq to identify regions of active transcription in human PDAC tissues. We identified SE regions containing SE-associated genes computationally for both normal and tumor samples (**Figure 1A**). We performed hierarchical clustering of the SE-associated genes that were enriched in tumor and normal tissues (**Figure S1A and Table S7**). We used Genomic Regions Enrichment of Annotations Tool (GREAT [22]) to determine which pathways were enriched in tumor versus normal-adjacent tissues.

**Figure 1.**
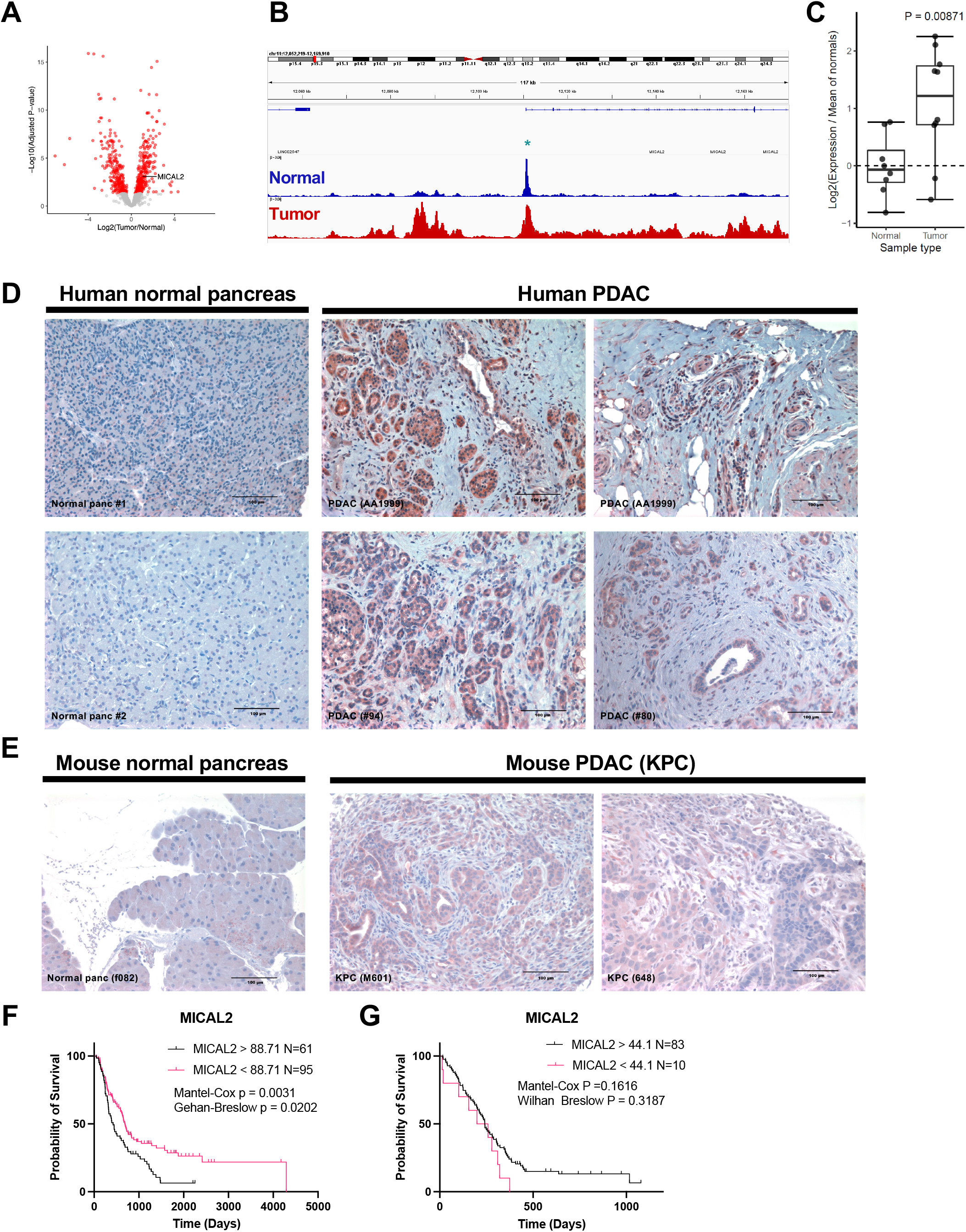
H3K27ac ChIP-Seq in human PDAC identifies MICAL2 as a SE associated gene. **A.** Differentially expressed SE-associated genes in tumor compared to normal tissue. MICAL2 gene is annotated. The red color denotes genes enriched with an adjusted p-value less than 0.05. **B.** H3K27ac ChIPseq occupancy upstream and within the MICAL2 loci in aggregated normal and tumor samples. The star denotes the MICAL2 gene start site. **C.** qPCR of MICAL2 using RNA extracted from the same patient samples used for ChIPseq. **D.** IHC for MICAL2 in normal pancreas and PDAC human tissues. **E.** IHC for MICAL2 in normal pancreas and KPC-derived PDAC mouse tissues. **F-G.** Survival analysis of PDAC patients segregated by MICAL2 expression (High vs Low) in two datasets: PanCuRx (**F**) and COMPASS (**G**).

Overall, we found that immune, KRAS and epithelial to mesenchymal transition (EMT) pathways were upregulated in tumor samples, while pancreatic endocrine genes and metabolic pathways were correlated with normal tissues (**Figure S1B**). Among the tumor-specific SE-associated genes we found leukemia inhibitory factor (Log Fold change 1.20; adjusted p value 6.20*10^-5^) which was previously identified as a potential therapeutic target in PDAC and is a target of ongoing clinical trials (13).

We next sought to identify SE-associated genes that were upregulated in PDAC, and which encoded proteins with the potential to be targeted by small molecules or antibody-based therapeutics. One of the top hits in our analysis was MICAL2, a member of the Molecule Interacting with CasL (MICAL) family. MICAL2 contains several functional protein domains including its flavin adenine dinucleotide enzymatic domain. MICAL2 was significantly enriched (Log Fold change 1.07; adjusted p value 5.39*10^-4^) in tumor samples compared to normal (**Figure 1A-B**). Furthermore, quantitative PCR analysis revealed that MICAL2 mRNA was enriched (p value 0.009) in tumor samples suggesting that the increased H3K27ac at the promoter and coding region does lead to increased transcriptional activation of the *MICAL2* loci (**Fig 1C**). We next assessed the enrichment of transcription factor binding motifs using GSEA and we found significant enrichment of development and inflammation-promoting transcription factor motifs such as HOX, NFKB and STAT1 (**Figure S1C**). Interestingly, SRF motifs were the most prevalent in tumor samples and MICAL2 indirectly promotes SRF transcription activity. Transcription factor motifs associated with normal pancreas development such as PAX and GATA genes were decreased in tumor tissues.

Investigation of TCGA solid cancers datasets revealed that PDAC is the 4^th^ highest expressor of MICAL2 (**Figure S1D**). Furthermore, using the TNMplot dataset [26] we determined that MICAL2 expression was significantly enriched in PDAC primary tumors (2.52 fold change; Dunn Test p value 4.60*10^-21^) and metastases (1.56 fold change; Dunn Test p value 2.58*10^-3^) compared to normal pancreatic tissues (**Figure S1E**). To ascertain if the higher level of MICAL2 transcription resulted in greater MICAL2 protein levels within PDAC tumors, we performed immunohistochemistry (IHC) on human normal pancreas and PDAC tissues. We found a striking tumor-specific increase in MICAL2 staining across 4 patient tumors (**Figure 1D**). We next observed that high MICAL2 protein levels and tumor-specificity are preserved in the commonly used Kras^LSL-G12D^; TP53^LSL-R172H^; and PDX1-cre (KPC) genetically engineered mouse model (**Figure 1E**) [27]. In addition, we found that MICAL2 was significantly overexpressed in murine KC and KPC organoids (**Figure S1F**) [28]. We also investigated the expression of MICAL2 across commonly used PDAC cell lines and found that most lines express MICAL2; ASPC1 is a high MICAL2 expressor while BxPc3 and MIAPaCa2 express low to no MICAL2 likely recapitulating the heterogeneity of PDAC (**Figure S1G**).

Finally, we investigated the association of MICAL2 expression with outcomes in PDAC patients who were eligible for surgical resection (PanCuRx [29]) and in patients with advanced disease (COMPASS [30]). We found that high MICAL2 expression correlated with worse outcome in the surgical cohort but was not prognostic in the advanced cohort (**Figures 1F and G**). This suggested a possible role for MICAL2 in the progression from primary to advanced disease. Overall, we found that PDAC has a distinct landscape of SE-associated genes that are linked with known PDAC biology. We found MICAL2 among these genes and determined that MICAL2 transcription is higher in PDAC compared to normal adjacent tissues both in our study as well as in other independent datasets and that this is associated with increased expression at the protein level. Importantly high MICAL2 expression is associated with a poorer prognosis in patients whose tumors were surgically removed indicating expressing high MICAL2 may mark tumors at increased recurrence and progression.

### MICAL2 expression is associated with KRAS and EMT signaling pathways

As MICAL2 is known to canonically regulate MRTF/SRF activity, we first sought to determine if this was occurring in the setting of PDAC. We checked the expression of common SRF target genes by qPCR following constitutive shRNA KD of MICAL2 in ASPC1 cells and found that many of the MRTF/SRF target genes were downregulated as we expected (**Figure S2A**). RhoA expression was dramatically reduced in MICAL2 KD cells, suggesting that MICAL2 may regulate a key promoter and target of SRF signaling. Notably, constitutive shRNA KD of MICAL2 leads to a decrease of MRTF-A and -B expression suggesting that prolong loss of MICAL2 activity depresses SRF transcription (**Figures S2A**). Interestingly, in ASPC1 and KPC46 cells, siRNA KD of MICAL2 led to the small increase in MRTF-A and -B expression which may reflect acute compensation (**Figure S2B-C**). To further understand the impact of MICAL2 on pancreatic cancer cell biology we performed RNA sequencing. Using differential gene expression analysis comparing siRNA targeting MICAL2 and scramble control, we found as expected that MICAL2 was the most significantly repressed gene (**Figure 2A and Table S8**). To investigate pathways likely to be regulated by MICAL2, we performed gene set enrichment analysis (GSEA). Interestingly, we found that KRAS signaling pathways were dramatically reduced in ASPC1 cells lacking MICAL2 (**Figure 2B and 2C**). Additional pro-survival pathways were lost in the MICAL2 KD cells such as TNFa and HIF1a signaling suggesting that MICAL2 may act as a proto-oncogene in PDAC. Importantly, we found that epithelial-to-mesenchymal transition (EMT) signaling was also significantly reduced in MICAL2 KD cells (**Figure 2C**). As an orthogonal pathway analysis method, we identified the cell states, elucidated as transcriptional activity downstream of *KRAS* activation via the OncoGPS methodology [31], that were associated with MICAL2 expression in other tumor models of the Cancer Cell Line Encyclopedia (CCLE) [32]. Similar to GSEA, we found that the EMT state was associated with higher MICAL2 gene expression in the CCLE (**Figure 2D**). To further investigate the EMT phenotype, we used a murine PDAC organoid line derived from a KPC liver metastasis, KPC46, which has mesenchymal features and has a high propensity for metastasis (**Figure S2C**). In MICAL2 KD cells, we observed mesenchymal to epithelial changes, marked by highly uniform, polarized, compact epithelial cell structures (**Figure 2E**). Overall, these experiments revealed that MICAL2 drives MRTF/SRF transcription, EMT, and pro-oncogenic pathways in both human and mouse models of PDAC.

**Figure 2.**
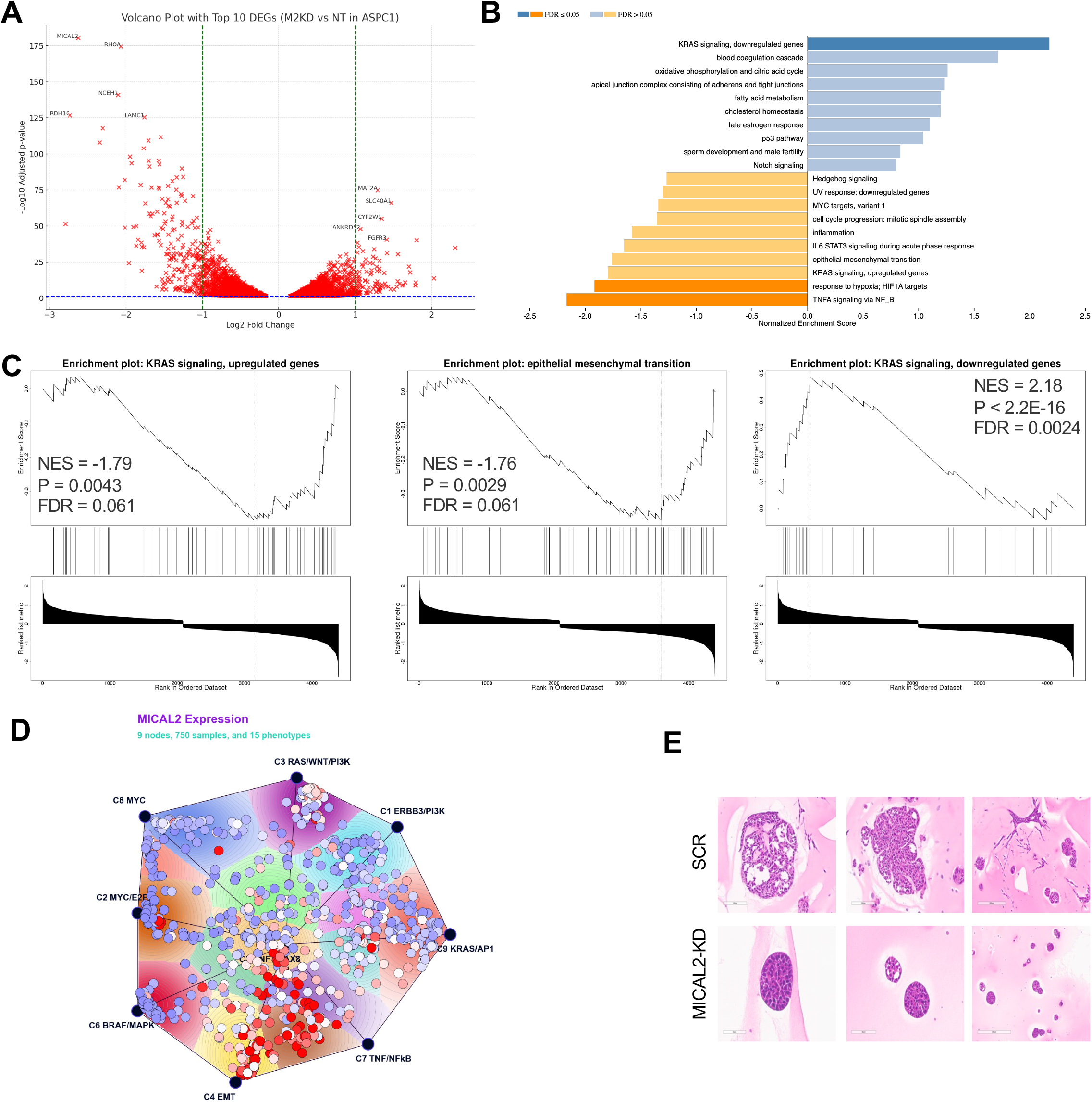
MICAL2 promotes a KRAS and EMT phenotype in PDAC cells. **A.** Differentially expressed genes in AsPC1 cells with MICAL2-KD (M2KD) compared to scramble control (NT), top 5 up- and top 5 down-regulated genes are annotated. **B.** GSEA Hallmark pathways enriched in AsPC1 cells with MICAL2-KD (blue) compared to pathways enriched in control AsPC1 cells (orange). **C.** GSEA enrichment plots indicating normalized enrichment score (NES), adjusted p value (P) and false discovery rate (FDR). **D.** OncoGPS plot of CCLE cell lines showing MICAL2 expression. Cell lines with high MICAL2 expression are indicated by red dots, and low expression by blue dots. **E.** H&E stain of mouse PDAC organoids derived from control KPC46 cells (SCR) and MICAL2-KD (MICAL2).

### MICAL2 promotes KRAS signaling

Our transcriptomic analyses indicated that KRAS signaling is altered in MICAL2-deficient PDAC cells. Therefore, we evaluated the activation of PI3K and MAPK signaling cascade in cells with loss and gain of function of MICAL2. Since MICAL2 loss leads to altered expression of the SRF co-activators MRTF-A and MRTF-B, we also investigated the effect of KD of these two genes. We generated human and mouse models silencing MICAL2, MRTF-A, and MRTF-B (**Figure S3A and S3B**). For gain of function studies, we overexpressed (OE) MICAL2 in BxPc3 human PDAC cells as BxPc3 does not endogenously express MICAL2 (**Figure S3C and S1G**). In the human PDAC cell line, AsPC1, loss of MICAL2 led to a marked decrease in p-AKT, and a minor decrease in p- ERK1/2 (**Figure 3A**). The negative cell cycle regulator P27 was dramatically increased in MICAL2 KD cells suggesting a possible mechanism for decreased cell proliferation. The siRNA-mediated silencing of MRTF-B phenocopied the p-AKT and p-ERK1/2 decrease observed in MICAL2 silenced cells. Notably, MICAL2 KD led to a partial loss of MRTF-B protein while the RNA was not decreased (**Figure S2A**). Interestingly, the loss of MRTF-A did not recapitulate this phenotype and neither MRTFs’ loss of function phenocopied the P27 increase. We next evaluated the loss of MICAL2 and MRTFs in KPC46 cells. We found that mouse PDAC cells recapitulated the decrease of p-AKT and pERK1/2 in both MICAL2 and MRTF-B silenced cells, however, P27 was unchanged in this cell line (**Figure 3B**). When we examined BxPc3 cells modified to express MICAL2, we found that p-AKT, p-ERK, and P27 expression were increased (**Figure 3C**). These results demonstrate that MICAL2 and MRTF-B deficient cells have reduced phosphorylation of PI3K and MAPK consistent with reduced KRAS signaling in these cells. To further assess the impact of MICAL2 on KRAS signaling, we examined the effects of MICAL2 KD on macropinocytosis, an extracellular nutrient scavenging process driven by KRAS. We observed a dramatic reduction in macropinocytosis in MICAL2 deficient cells which is also consistent with a decrease in KRAS signaling activity (**Figure 3D-E**). In sum, loss, and gain of function studies revealed that MICAL2 promotes signaling through KRAS as manifest by phosphorylation of MAPK and PI3K and macropinocytosis. These observed biochemical results were thus consistent with our pathway analyses of the cellular transcriptome.

**Figure 3.**
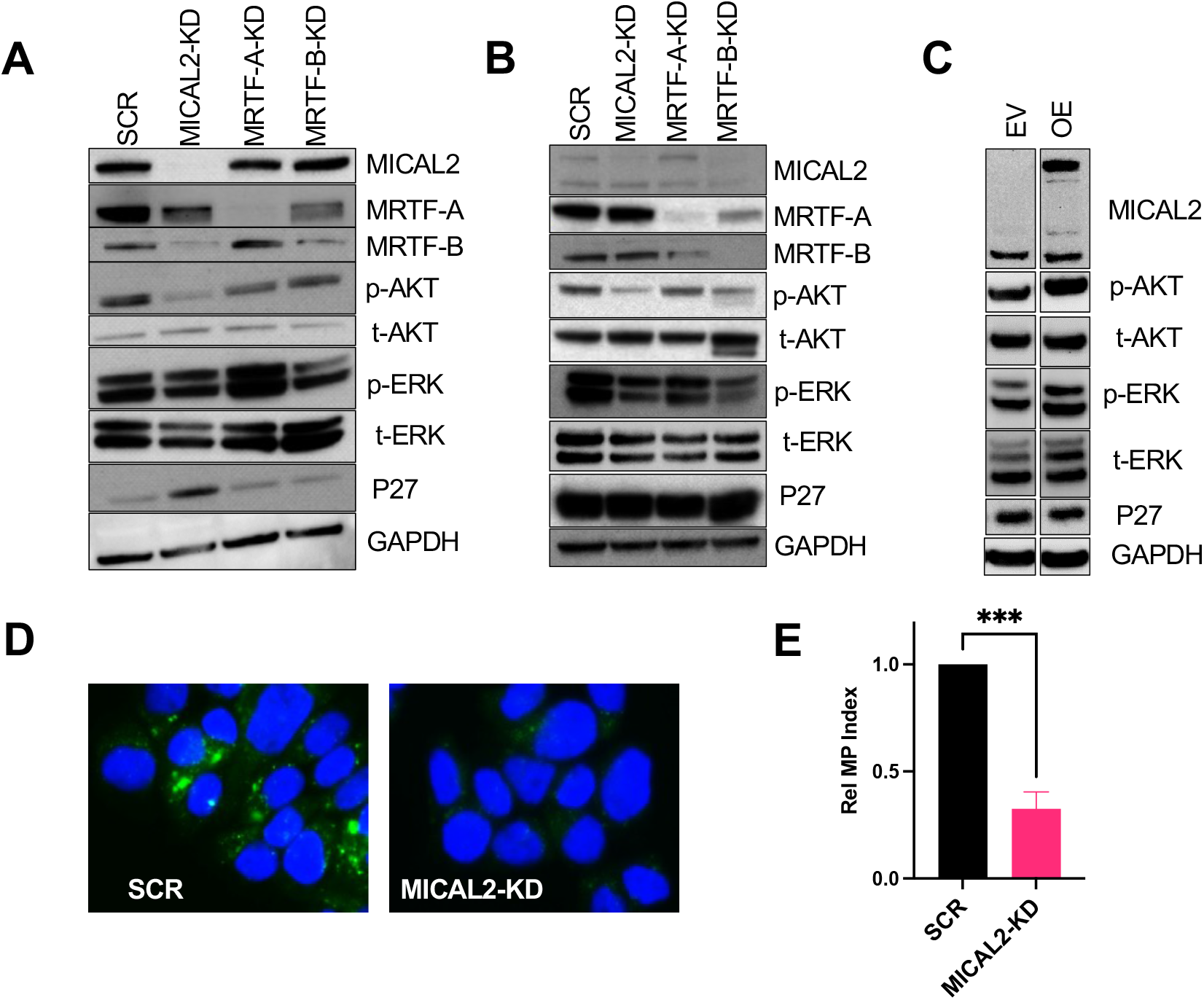
MICAL2 promotes KRAS signaling through AKT and ERK ex vivo. **A-B.** Immunoblot analysis of AsPC1 (**A**) and KPC46 (**B**) cells treated with SCR, MICAL2, MRTF-A and MRTF-B siRNAs at 72 hrs. **C**. Immunoblot analysis of BxPc3 cells expressing empty vector (EV) or MICAL2-overexpression vector (OE) at 72 hrs. **D.** Representative immunofluorescent images of AsPC1 cells transfected with siRNA control (SCR) or MICAL2. DAPI (blue) marks cell nuclei, and FITC-conjugated dextran (green) is used to label macropinosomes. **E.** Quantitation of the relative macropinosome index from D.

### MICAL2 promotes PDAC cell proliferation and migration

We next sought to investigate how MICAL2 loss and gain of function would impact oncogenic phenotypes in PDAC cells. Since MICAL2 appeared to drive EMT, we first measured the motility of MICAL2, MRTF-A, and MRTF-B deficient cells. We found that migration in both ASPC1 and KPC46 was reduced after MICAL2 KD (**Figure 4A and B**, **Figure S4A and B**). We also found that MRTF-A and -B KD decreased migration in the human ASPC1 cells, while in mouse KPC46 only MRTF-A KD resulted in reduced migration. Conversely, in a gain-of-function experiment, BxPc3 cells overexpressing MICAL2 had increased migration compared to empty vector control cells (**Figure 4C; Figure S4C**). Using an invasion assay, we further determined that KPC46 cells with loss of MICAL2 were less capable of invading a gelatin matrix (**Figure S4D**).

**Figure 4.**
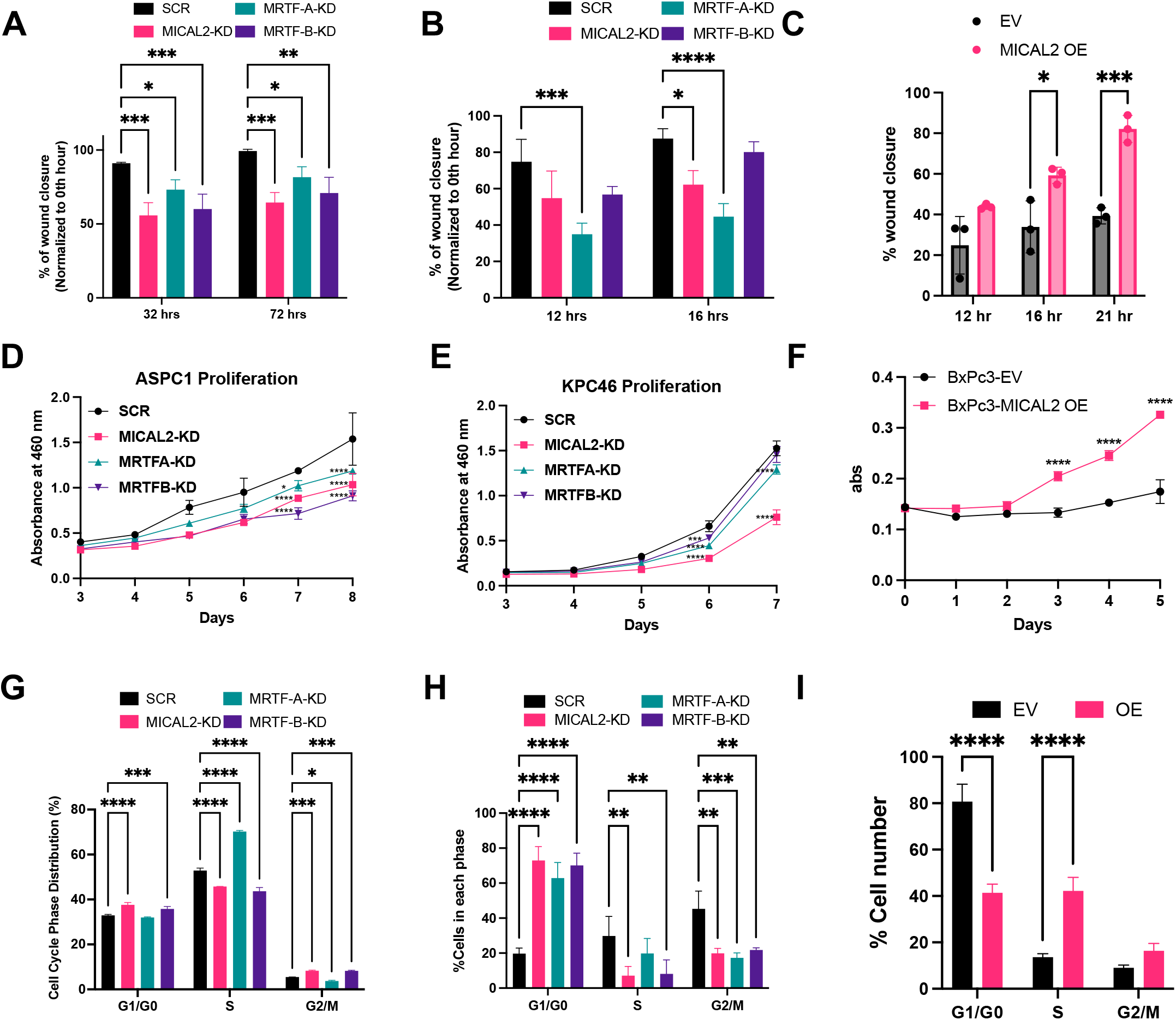
MICAL2 drives cancer cell migration and proliferation in vitro. **A-B.** Quantification of wound healing assay of AsPC1 (**A**), and KPC46 (**B**) transfected with SCR, MICAL2, MRTF-A and MRTF-B siRNAs at the time points indicated. **C.** Quantification of wound healing assay of BxPc3 cells expressing empty vector (EV) or MICAL2-overexpression vector (OE) at the time points indicated. **D-E.** Proliferation assay of AsPC1 (**D**), and KPC46 (**E**) transfected with SCR, MICAL2, MRTF-A and MRTF-B siRNAs at the time points indicated. **F.** Proliferation assay of BxPc3 cells expressing empty vector (EV) or MICAL2-overexpression vector (OE) at the time points indicated. **G-I.** Cell cycle analysis of AsPC1 (G), KPC46 (**H**) transfected with SCR, MICAL2, MRTF-A and MRTF-B siRNAs at 72 hrs, and BxPc3 (**I**) cells expressing empty vector (EV) or MICAL2-overexpression vector (OE).

We next assessed the impact of MICAL2 on cell proliferation. We found that MICAL2 KD led to a significant decrease in proliferation in ASPC1 and KPC46 cells, while OE of MICAL2 in BxPc3 led to an increase in proliferation rate compared to control cells (**Figures 4D-F**). Interestingly, KD of either MRTF in the human or mouse PDAC cells led to only a partial decrease in the cell proliferation rate (**Figures 4D-E**). To better understand which part of the cell cycle was impacted by MICAL2 expression, we used flow cytometry to examine cell cycle progression. In both AsPC1 and KPC46, MICAL2 and MRTF-B deficient cells had a significant shift toward arrest in the G0/G1 phase and concomitant reduction in the proportion of cells in the S phase and G2/M (**Figure 4G and H**). MRTF-A-depleted KPC46 cells had a cell cycle profile but AsPC1 cells lacking MRTF-A surprisingly had a block in S phase rather than G0/G1 hinting at differences between models and possibly between human and mouse PDAC cells. Conversely, BxPc3 MICAL2-OE cells progressed faster through G0/G1 and had an increased S phase proportion compared to the control indicating that the increase in MICAL2 expression was sufficient to increase cell division (**Figure 4I**).

Overall, these experiments show that MICAL2 and MRTF-A/B expression promote cell migration invasion, and proliferation. Further, these findings are consistent with the RNAseq and biochemical results we observed after the genomic knockdown of MICAL2 in PDAC cells.

### MICAL2 and MRTF-B promote heterotopic and orthotopic growth *in vivo*

When we silenced MICAL2 *in vitro* we observed reduced cell proliferation, migration, invasion, and a reversal of the EMT phenotype. Therefore, we next sought to determine how the loss of MICAL2 and MRTFs impacted tumorigenesis initially using heterotopic subcutaneous mouse transplant models. We first transplanted AsPC1 cells with constitutive KD of MICAL2 or MRTF-A/B into immunodeficient NSG mice. We observed a dramatic decrease in tumor size when MICAL2 and MRTF-B were lost but no significant differences after MRTF-A silencing (**Figure 5A- B**). Transplantation of KPC46 cells in syngeneic mice recapitulated the results observed with human PDAC cells, though with an even more profound reduction in tumor formation (**Figure 5C- D**). In one experiment, no tumors formed from the MICAL KD cells while in a repeat experiment, 1/6 tumors didn’t form, and the remaining tumors were markedly growth inhibited relative to control. Conversely, BxPc3 MICAL2-OE cells grew larger than control cells when injected into the flank of NSG animals (**Figure 5E-F**).

**Figure 5.**
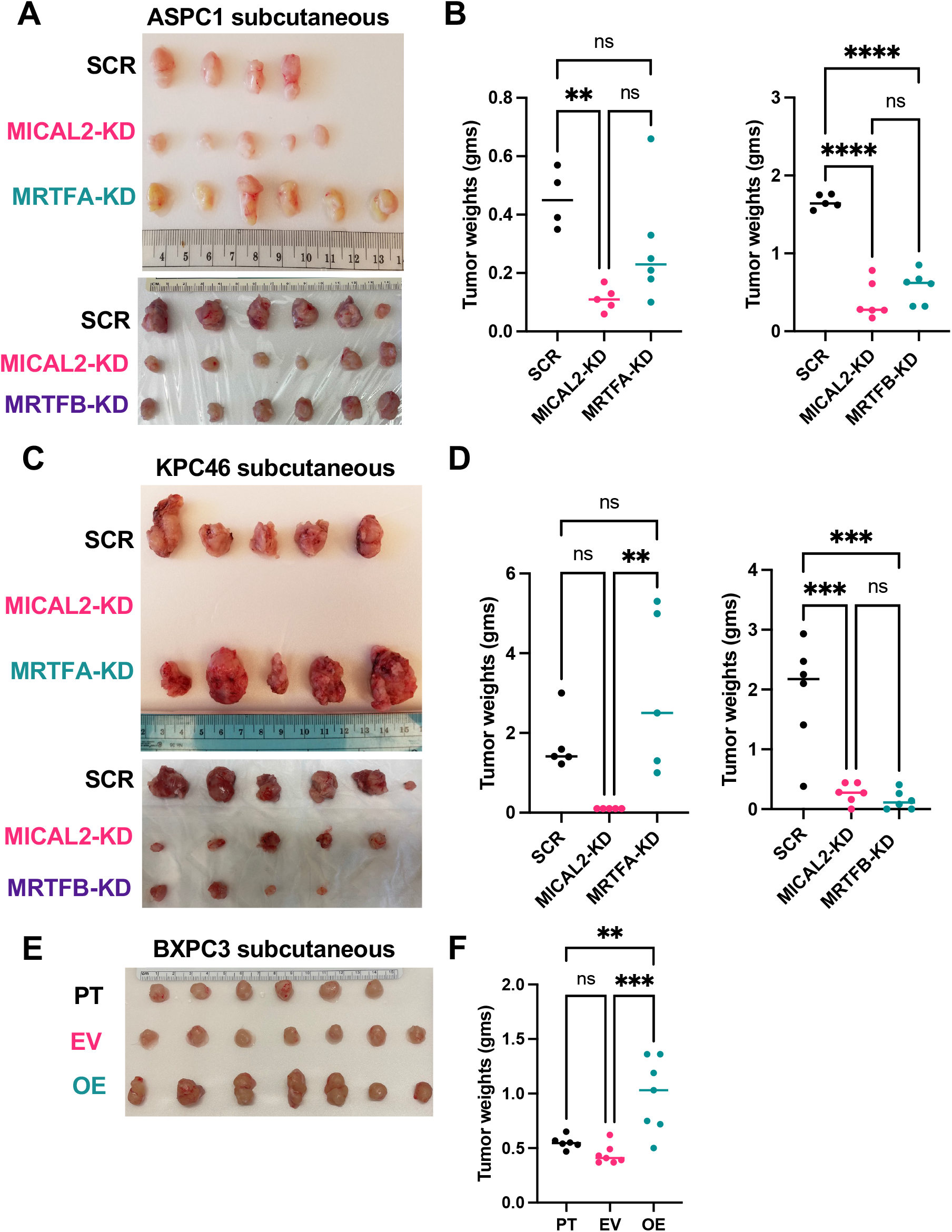
MICAL2 and MRTF-B promote heterotopic tumor growth. **A.** Representative images of subcutaneous AsPC1 tumors grown in immunocompromised mice. AsPC1 cells express shRNA vectors to silence MICAL2, MRTF-A, and MRTF-B. **B.** Weight quantification of the AsPC1 tumors shown in A. **C.** Representative images of subcutaneous KPC46 tumors grown in syngeneic mice. KPC46 cells express shRNA vectors to silence MICAL2, MRTF-A and MRTF-B. **D.** Weight quantification of the KPC46 tumors shown in C. **E.** Representative images of subcutaneous BxPc3 tumors grown in immunocompromised mice. BxPc3 cells express EV or MICAL2-OE vectors. **F.** Weight quantification of BxPc3 tumors shown in E.

We next sought to investigate how MICAL2 and MRTFs impact orthotopic tumor growth by implanting cells into the pancreatic tail. We found that only the mice implanted with AsPC1 MICAL2-KD cells, not the MRTF-KDs had a significantly decreased tumor burden, whereas in the KPC46 model, both the MICAL2 and MRTF-B silenced cells had reduced *in vivo* growth compared to scramble control (**Figure 6 A-D**). MRTF-A again had no impact on tumor growth. To evaluate if MRTF-B could compensate for the loss of MICAL2, we used an SRF reporter assay in KPC46 MICAL2 KD cells with and without the overexpression of MRTF-B. We found that, as expected, SRF signaling was downregulated in the KPC46 MICAL2 KD cells, but this effect was rescued by the OE of MRTF-B, demonstrating that MICAL2-dependent SRF signaling in PDAC is likely driven by modulating levels of MRTF-B (**Figure S5**). In summary, the expression of MICAL2 and MRTF-B in PDAC cells promoted tumor growth in both heterotopic and orthotopic locations whereas MRTF-A expression did not impact tumor growth. These results are consistent with our *in vitro* findings that MICAL2 promotes PDAC cell proliferation and suggest that MRTF-A and -B play distinct roles in promoting pancreatic tumor growth.

**Figure 6.**
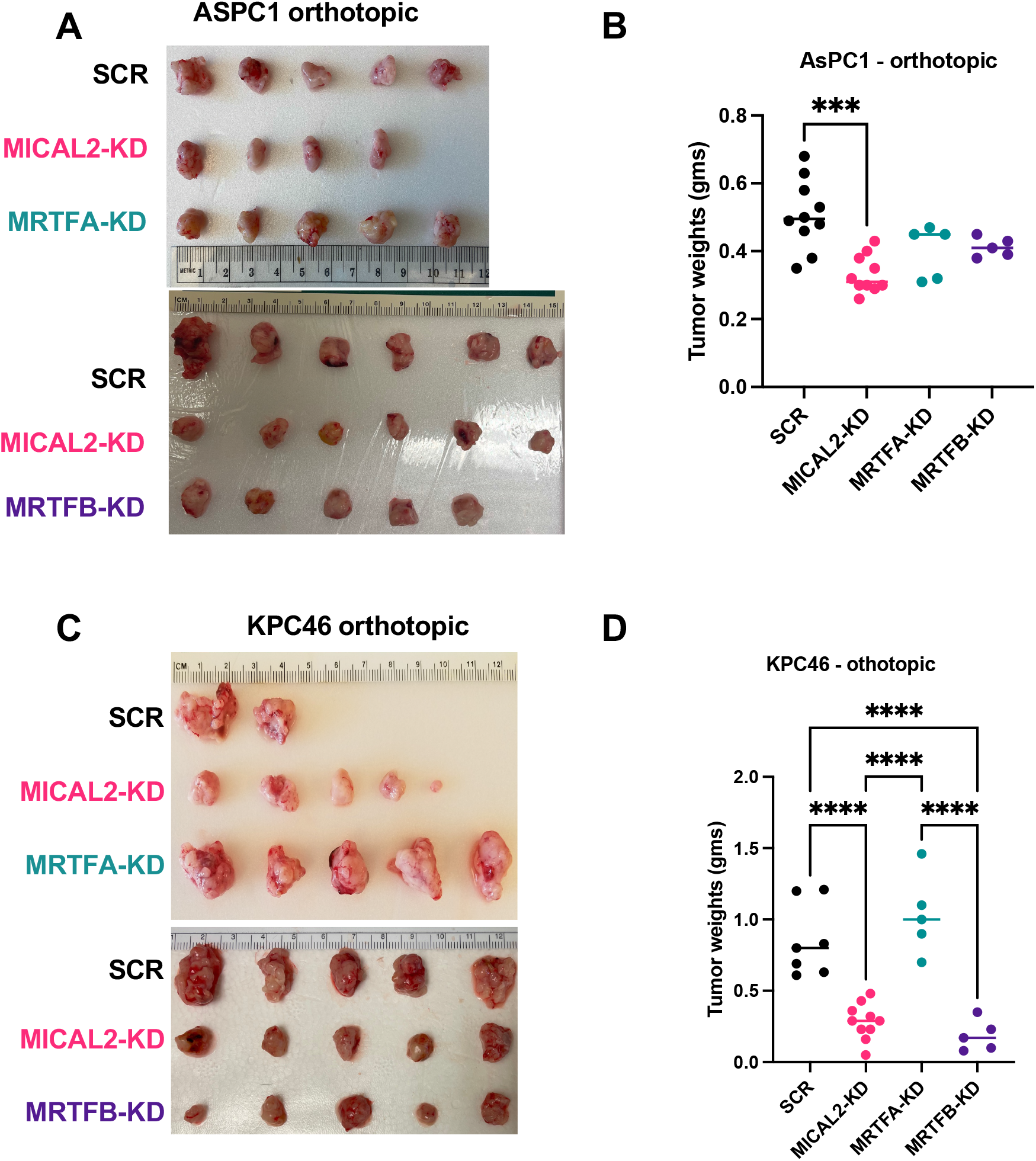
MICAL2 and MRTF-B promote orthotopic tumor growth. **A.** Representative images of orthotopic AsPC1 tumors grown in immunocompromised mice. AsPC1 cells express shRNA vectors to silence MICAL2, MRTF-A and MRTF-B. **B.** Weight quantification of the AsPC1 tumors shown in A. **C.** Representative images of orthotopic KPC46 tumors grown in syngeneic mice. KPC46 cells express shRNA vectors to silence MICAL2, MRTF-A and MRTF-B. **D.** Weight quantification of the KPC46 tumors shown in C.

### MICAL2 promotes metastasis in mice

To determine how MICAL2 and MRTFs expression impact the competency of PDAC cells to metastasize to the liver, we injected PDAC cells into the spleen of mice. We first injected KPC46 cells with KD of MICAL2, MRTF-A, and MRTF-B (**Figure 7A-B**). Grossly, we observed a dramatic decrease in liver metastatic burden in the MICAL2, MRTF-A and MRTF-B KD models compared to scramble control cells. Histologically, we found only small, and often merely microscopically detectable liver metastases in the KD models compared to extensive gross metastatic disease when the control cells were injected (**Figure 7C**).

**Figure 7.**
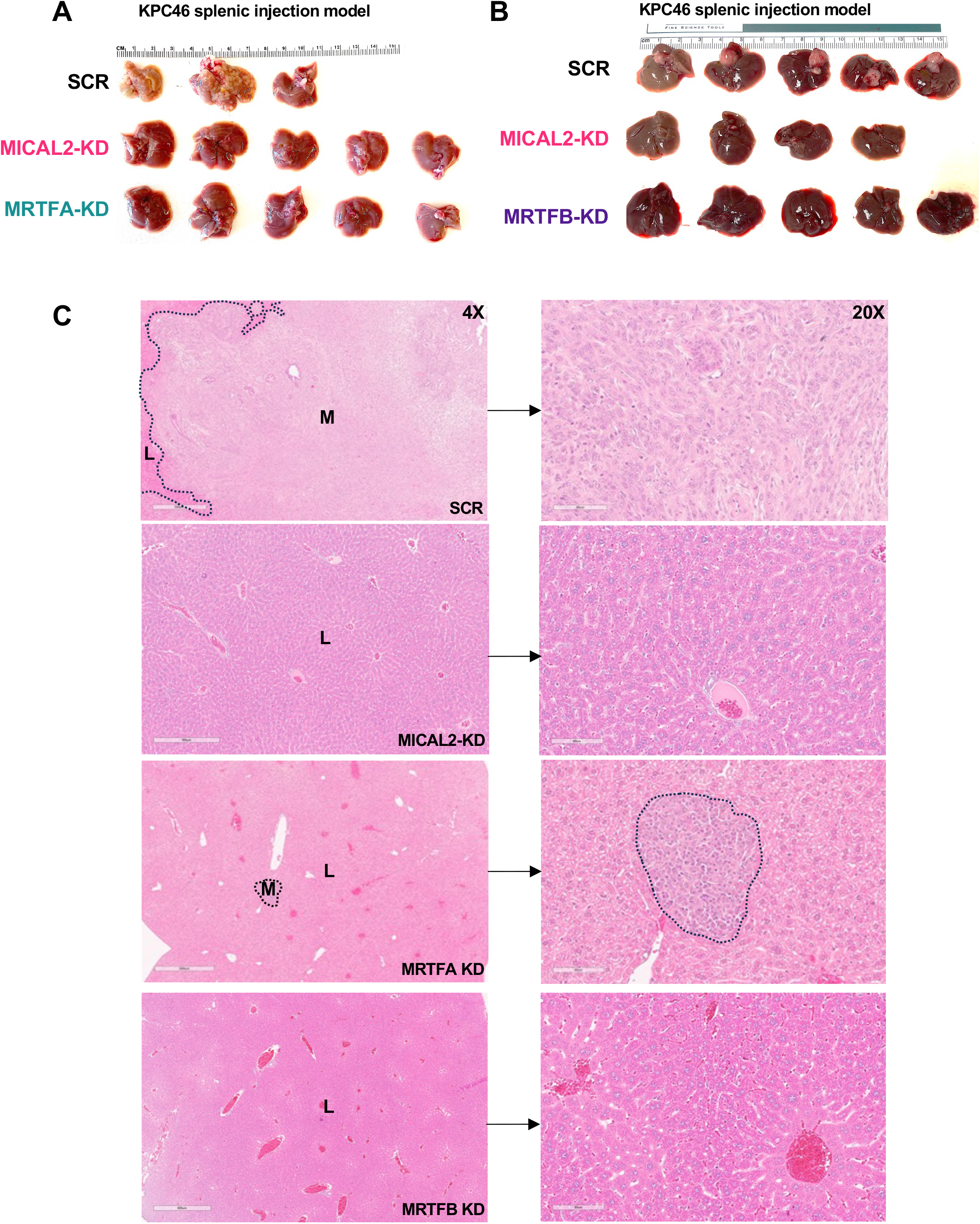
MICAL2, MRTF-A and MRTF-B promote metastatic spread *in vivo*. **A.** Representative images of liver metastatic burden after splenic injection of KPC46 cell into syngeneic mice. KPC46 cells express shRNA vectors to silence MICAL2, MRTF-A. **B.** Representative images of liver metastatic burden after splenic injection of KPC46 cell into syngeneic mice. KPC46 cells express shRNA vectors to silence MICAL2, MRTF-B. **C.** Representative H&E images of livers shown in A-B. Normal liver tissue (L) and metastasis (M, dotted lines) are labeled.

We then injected BxPc3 cells overexpressing MICAL2 or vector control into the spleen of NSG mice. While we did not observe gross metastatic disease in either group, microscopic analysis revealed numerous metastatic foci in the livers of mice implanted with BxPc3 MICAL2 overexpressing cells while there was no metastatic disease detectable in the control group (**Figure S6A-B**). These results suggest that MICAL2, MRTF-A, and B promote liver metastasis of PDAC cells.

## DISCUSSION

There remains a critical need to identify better pharmacological targets that drive pancreatic cancer. While commonly mutated, effectively targeting mutations in KRAS, CDKN2A, SMAD4, and TP53 has not been possible until the recent development of KRAS inhibitors, and no drugs have yet gained approval for pancreatic cancer therapy. Importantly, these and other known genetic alterations do not sufficiently explain the vast tumor heterogeneity and unique biology of PDAC [33, 34]. Patient outcomes, including survival, associated with PDAC subtypes have been strongly linked to epigenomic alterations in oncogenes and tumor suppressors [35–38].

Super-enhancers, as marked by H3K27 acetylation, are clusters of enhancers with aberrantly high levels of transcription factor binding, which are central to driving the expression of genes that control cell identity and stimulate oncogenic transcription [4]. Certain environmental or patient-specific events during tumorigenesis likely modulate specific super-enhancers to influence target gene expression associated with PDAC phenotypic outcomes [38]. We hypothesized that characterizing the landscape of H3K27 acetylation in pancreatic cancer tissue would lead us to identify super-enhancer-associated genes critical in driving tumor progression and maintenance. Although there have been efforts to target global epigenetic changes [39–41] and thereby mitigate PDAC-promoting pathways, our study is one of the first to directly perform unbiased discovery using an epigenetic screen on primary tumor and normal pancreatic tissue.

Our characterization of super-enhancer-associated regions through H3K27ac ChIP-Seq identified multiple differentially acetylated regions of the genome between tumor and normal tissues. Among them included leukemia inhibitory factor, previously identified as a putative target in pancreatic cancer and a subject of ongoing clinical trials [42]. We chose to focus on the MICAL2 gene as it was one of the most differentially acetylated genes between PDAC and normal human pancreas. Furthermore, the enrichment of the SRF transcriptional motif in PDAC associated SE also provided strong rationale to pursue MICAL2. MICAL2 is a flavin monooxygenase and families of flavo-proteins have been successfully targeted in human disease, i.e. monoamine oxidases (MAO) inhibitors [43] for depression and neurodegenerative disorders [44], as well as inhibitors of flavo-containing monooxygenases (FMOs) like methimazole [45] to target excess thyroid hormone synthesis. Hence, there is significant foundation to posit that it may be possible to inhibit the enzymatic domain of MICAL2 pharmacologically. MICAL2 canonical function is linked to transcriptional control of MRTF/SRF signaling. MRTF/SRF signaling has been shown to regulate transcription of genes that promote tumor EMT, fibrosis, and metastasis, which are hallmarks of PDAC biology [46]. In previous studies, MICAL2 has been primarily shown to promote EMT and migration in malignancies such as lung [11], gastric [12], and breast cancer [13] cell carcinoma. In addition, a single prior publication suggested that MRTF-A and B were oncogenic based on their modulation in PDAC cell lines [47]. Importantly, no prior studies have investigated the impact of MICAL2 on PDAC cells or studied its relationship to MRTF-A/B/SRF signaling in this disease. At the time of manuscript submission, using bioinformatic approaches Liu et al. reported that MICAL2 is overexpressed in PDAC and that its expression was associated with EMT and poor prognosis, findings entirely consistent with our own [48]. They further noted that MICAL2 is expressed in PDAC associated fibroblasts and an immunosuppressive microenvironment, findings that only serve to heighten our interest in MICAL2 as a putative therapeutic target.

After identifying MICAL2 as a super enhancer-associated gene, we confirmed its overexpression at the RNA and protein level in both human and mouse PDAC cells. *In vitro*, we found MICAL2 to be a driver of cell proliferation, cell cycle progression, matrix invasion, macropinocytosis and migration while *in vivo*, we determined that MICAL2 promotes tumor growth and metastasis in several model systems. We have shown that MICAL2 expression regulates MRTF-SRF transcriptional targets and that the MICAL2-regulated factors MRTF-A and -B also play a critical role in PDAC biology. These findings, for the first time to our knowledge, clearly implicate SRF transcription as an important driver of PDAC gene expression. Silencing of MICAL2 and MRTF- B lead to a marked decrease in tumor growth in both heterotopic and orthotopic mouse models, suggesting their important role in tumorigenesis. We found that MICAL2, and MRTF-A and B, promote PDAC metastasis. Our findings that link MICAL2-promoted tumor progression specifically to MRTF-B are novel, as is also the identification of a regulatory loop whereby, they reciprocally promote each other’s transcription. Yet, our study also reveals a clear distinction between the impact of MRTF-A and MRTF-B on PDAC tumorigenesis and progression, as MRTF-A promoted metastasis but not tumor growth, whereas MRTF-B was necessary for both metastasis and tumor growth demonstrating a differential role of MRTF isoforms in PDAC.

We made some unexpected and novel observations that further heighten our interest in MICAL2 as a putative target for pancreatic cancer therapy, namely our RNAseq data suggesting that loss of MICAL2 expression downregulates genes associated with KRAS signaling. We further found that loss of MICAL2 expression in vitro resulted in decreased ERK1/2 and AKT phosphorylation. Although ERK1/2 and PI3K are activated downstream of various signals, the impact of MICAL2 may be linked to its effects on KRAS. This hypothesis is bolstered by our findings that loss of MICAL2 expression significantly reduced macropinocytosis, a KRAS-driven nutrient scavenging activity. Given the central role of KRAS in pancreatic cancer biology, further studies of the mechanisms by which MICAL2 impacts KRAS signaling are of great interest. Interestingly, neither MTRF-A or -B loss consistently resulted in a profound decrease in ERK1/2 and AKT activation. This suggests that the link of MICAL2 to KRAS signaling may be independent of its role in promoting MRTF/SRF transcription.

Our study expands upon previous reports on the role of MICAL2 in cancer by providing a comprehensive analysis of its role in PDAC. It further reveals, for the first time, that MICAL2 impacts KRAS related gene expression as well as KRAS regulated macropinocytosis. However, many questions remain, most specifically related to the mechanisms by which MICAL2 promotes these oncogenic phenotypes and KRAS function. The canonical function of MICAL2 in promoting MRTF/SRF expression likely underpins many of these effects, however, it is also localized in the cytosolic protein with other functional domains that may play an important role in MICAL2 related phenotypes.

In conclusion, we have studied the epigenomic landscape of PDAC to identify MICAL2 as a unique driver of PDAC progression through its regulation of MRTF/SRF signaling and determined differential effects of MRTF-A versus B in promoting PDAC growth. MICAL2 and MRTF-SRF biology represent attractive novel and potentially tractable targets for PDAC therapy given their regulation of transcriptional programs fundamental to its tumor biology.

## ACKNOWLEDGEMENTS

The authors gratefully acknowledge support from The Lustgarten Foundation (AML, JM, PB), Pancreatic Cancer Action Network (AML), NIH grants CA 273973, CA274295 and CA 285115 (AML), Stand Up to Cancer (AML), and the Research for a Cure of Pancreatic Cancer fund (AML). We also thank the Biorepository and Tissue technology shared resource for Biospecimen collection which is supported by CCSG Grant P30CA23100. This study was conducted with the support of the Ontario Institute for Cancer Research (PanCuRx Translational Research Initiative) through funding provided by the Government of Ontario, the Wallace McCain Centre for Pancreatic Cancer supported by the Princess Margaret Cancer Foundation, the Terry Fox Research Institute, the Canadian Cancer Society Research Institute, and the Pancreatic Cancer Canada Foundation. The study was also supported by a charitable donation from the Canadian Friends of the Hebrew University (Alex U. Soyka). Steven Gallinger is the recipient of an Investigator Award from OICR. The study was also supported by NIH grants CA 207189 (CC), NIH grant U24CA220341 (JPM and AW), Cancer Center Support Grant P30 CA051008 (JPM), and NIH grant CA 257344 (AW).

---

**Supplementary Figure 1.**
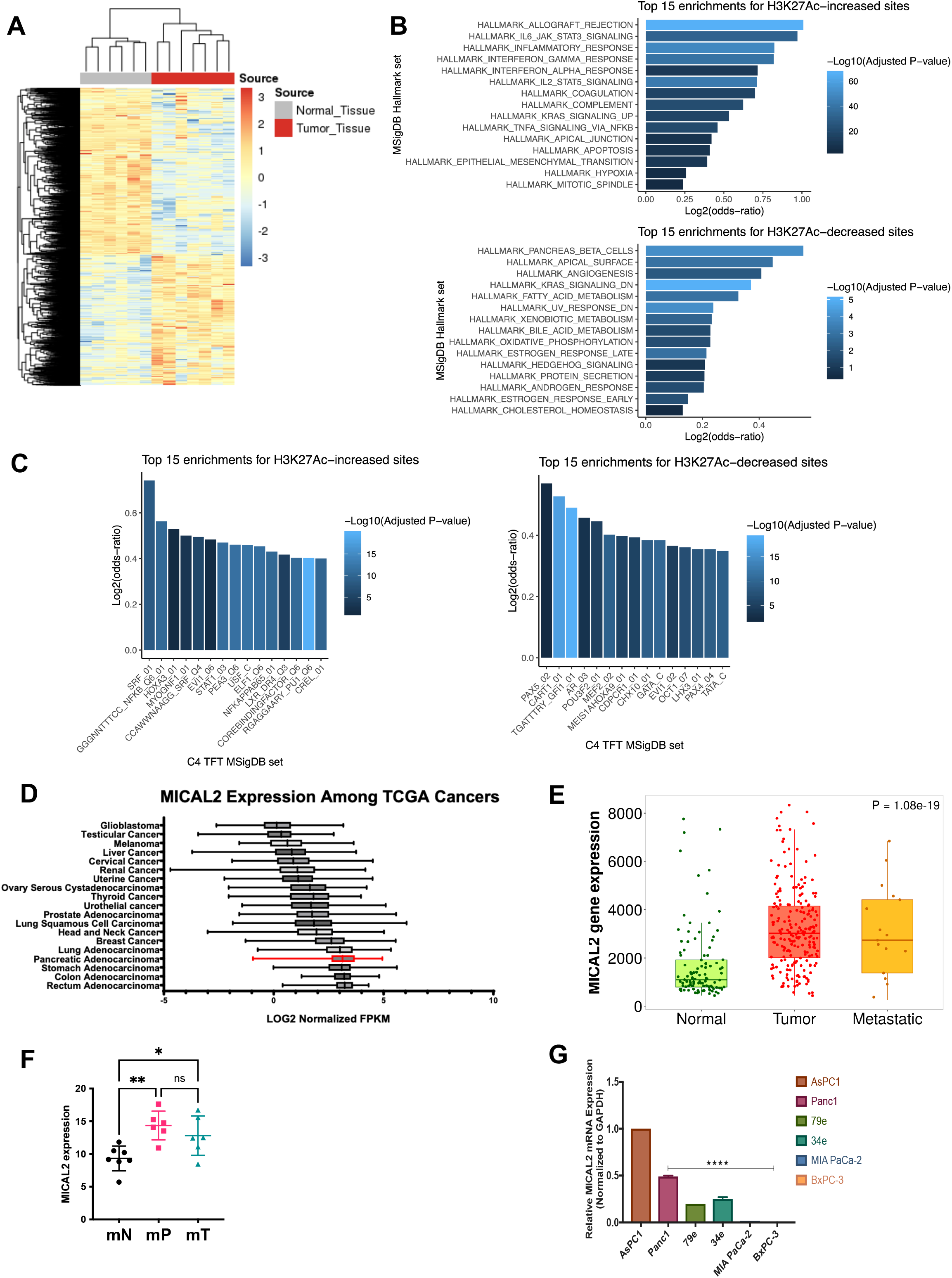
**A.** Heatmap of SE-associated genes in normal pancreas and PDAC tissues. **B.** Genes with increased or decreased H3K27Ac signals were subjected to GSEA analysis. **C.** Transcription factors motifs increased or decreased within H3K27Ac signals were subjected to GSEA analysis. **D.** MICAL2 expression in solid tumor TCGA datasets. Data are ranked by mean expression in each dataset from bottom (highest mean expression) to top (lowest mean expression). **E.** MICAL2 expression in normal pancreas, PDAC tumor and metastasis using the TNMplot dataset. **F.** MICAL2 expression by RNAseq in mouse organoids derived from normal pancreatic ducts (WT mice, mN), early PanIN lesions (KC mice, mP) and PDAC (KPC tumor bearing mice, mT). **G**. MICAL2 expression by qPCR in human PDAC cell lines.

**Supplementary Figure 2.**
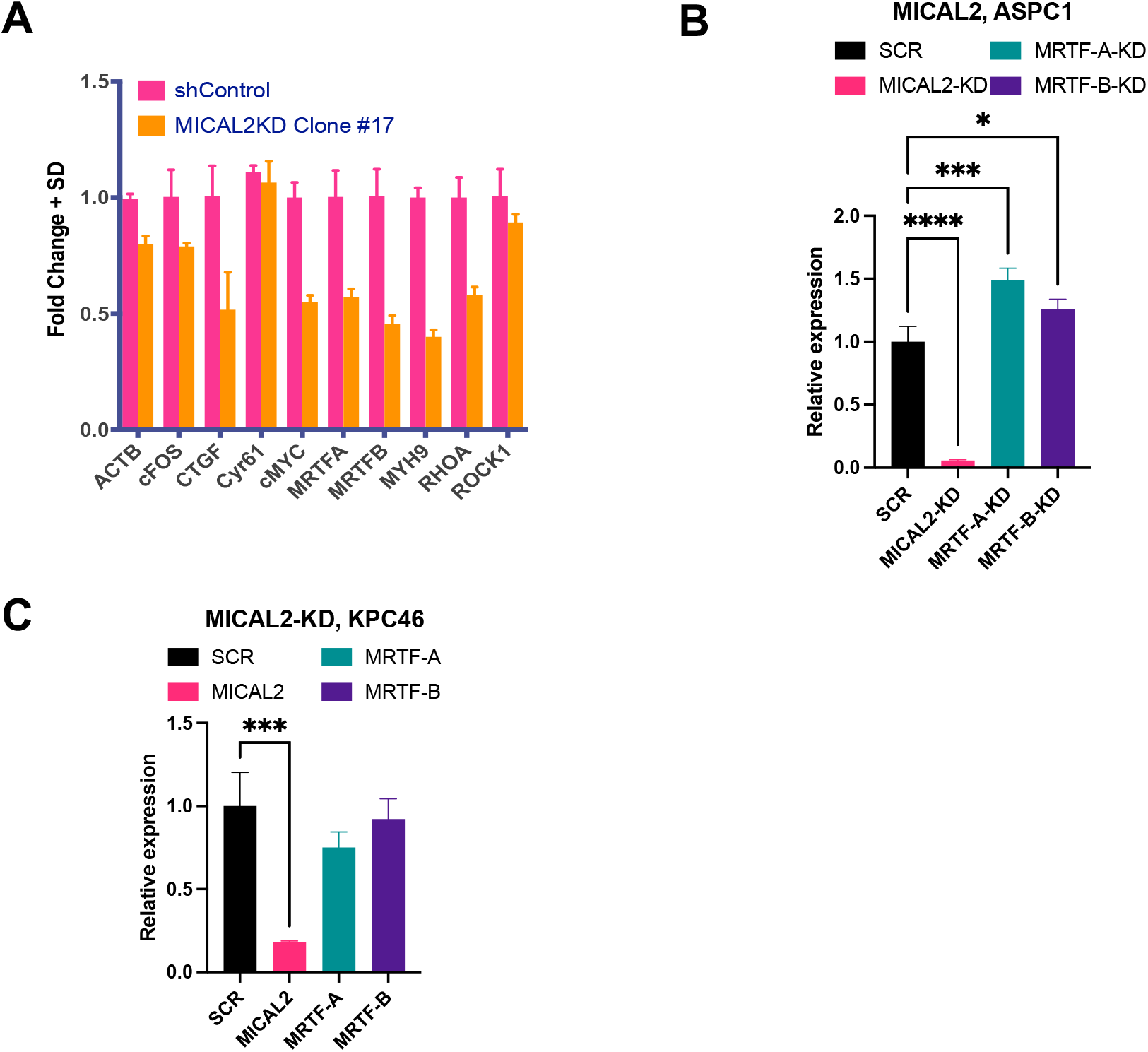
**A.** Normalized qPCR expression of SRF related genes in AsPC1 cells with scramble and MICAL2 shRNA expression. **B.** Normalized qPCR expression of MICAL2, MRTF-A, and MRTF-B in AsPC1 cells with siRNA KD of MICAL2. SCR denotes the scramble siRNA control. **C.** Normalized qPCR expression of MICAL2, MRTF-A and MRTF-B in KPC46 cells with siRNA KD of MICAL2. SCR denotes the scramble siRNA control.

**Supplementary Figure 3.**
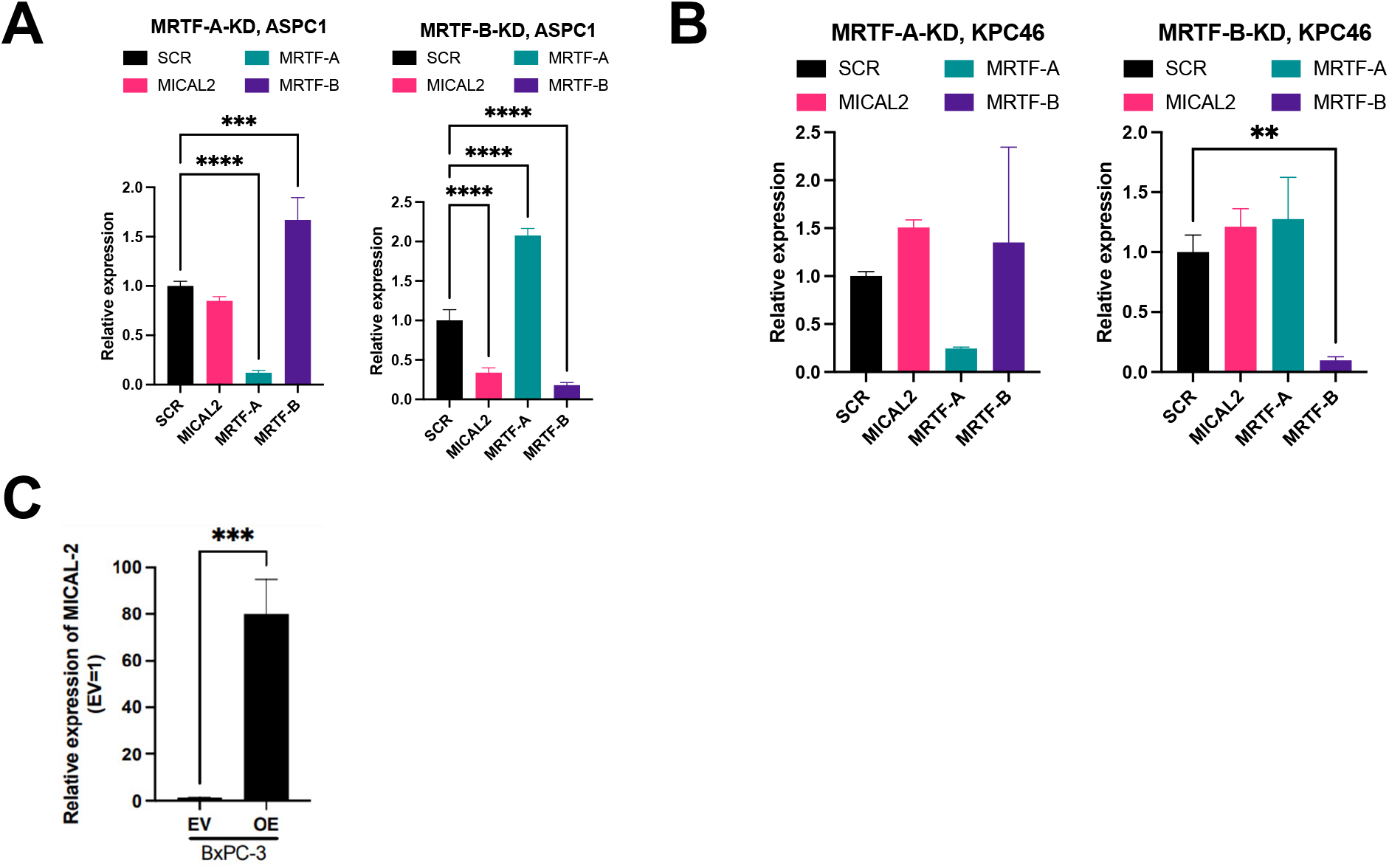
**A-B.** Normalized qPCR expression of MICAL2, MRTF-A and MRTF-B in AsPC1 (**A**) and KPC46 (**B**) cells with siRNA KD of MRTF-A and MRTF-B. SCR denotes the scramble siRNA control. **C.** Normalized qPCR expression of MICAL2 in BxPc3 cells expressing empty vector (EV) or MICAL2-overexpression vector (OE).

**Supplementary Figure 4.**
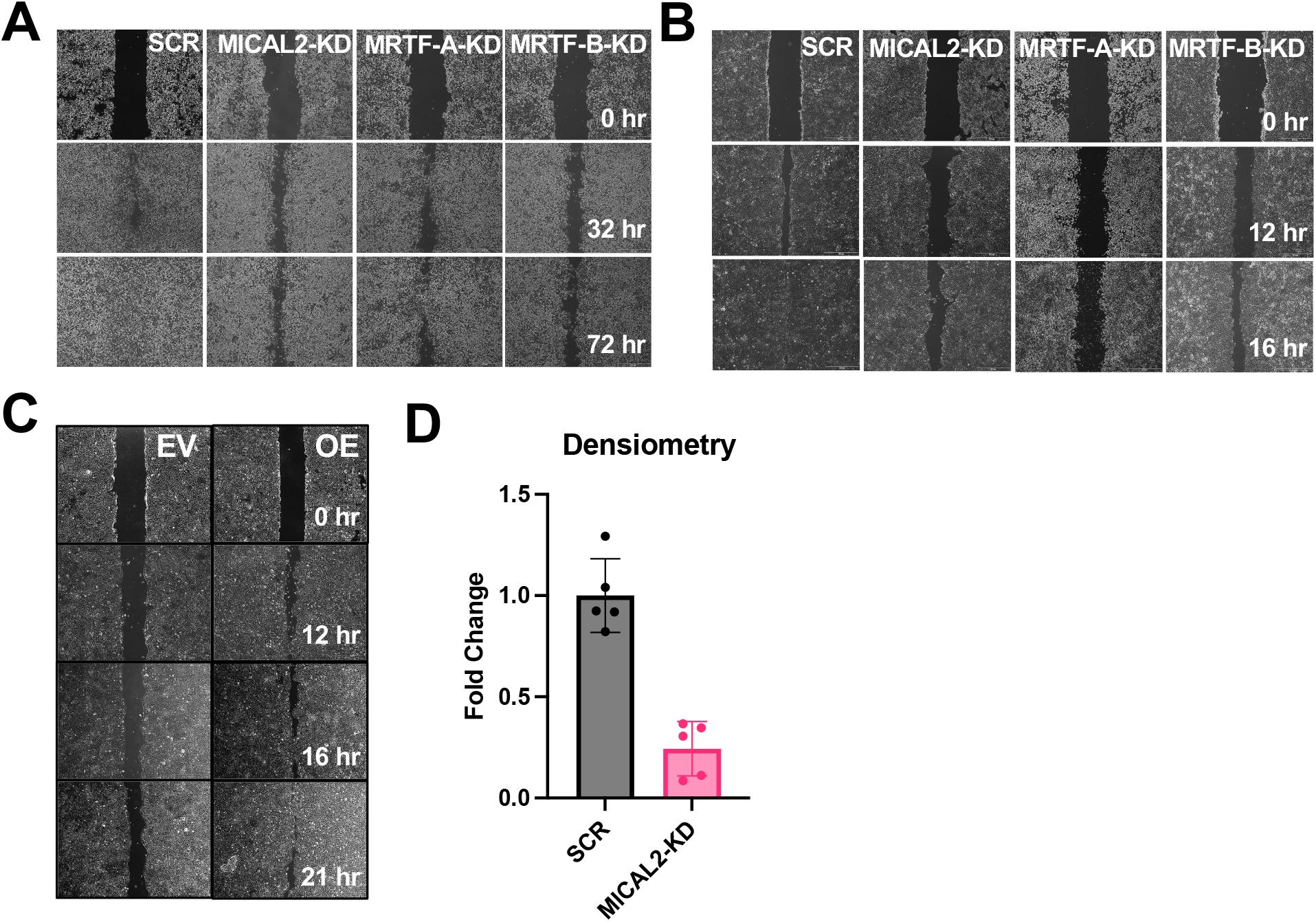
**A-C.** Representative images of wound healing assays quantified in Figure 4 for AsPC1 (**A**), KPC46 (**B**) and BxPc3 (**C**) cells. **D.** Densitometry analysis of gelatin matrix degradation/invasion by KPC46 cells transfected with control or MICAL2 targeted siRNA.

**Supplementary Figure 5.**
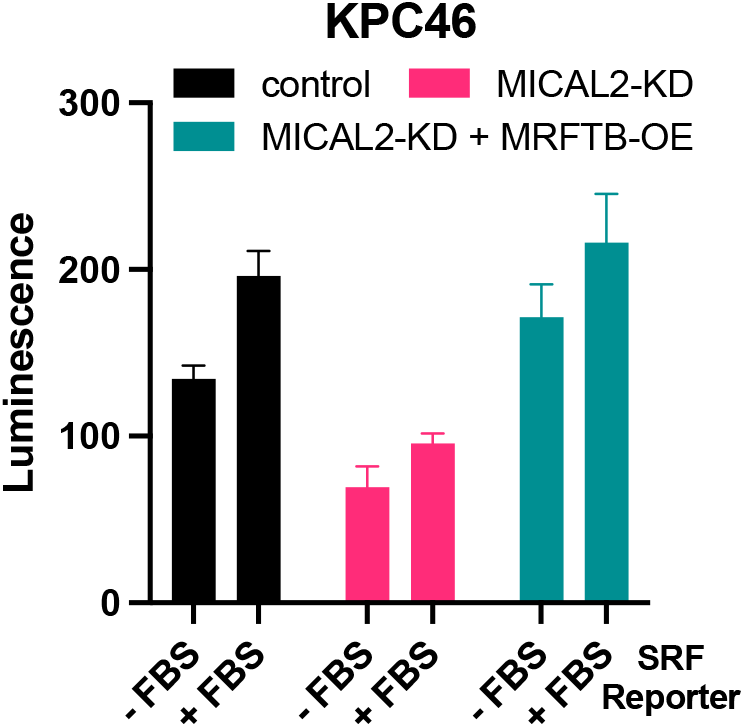
SRF reporter assay performed using KPC46 cells expressing shRNA vectors targeting MICAL2 or control. A secondary vector is overexpressing MRTF-B in the background of MICAL2-KD. SRF activity is induced by addition of FBS to the media.

**Supplementary Figure 6.**
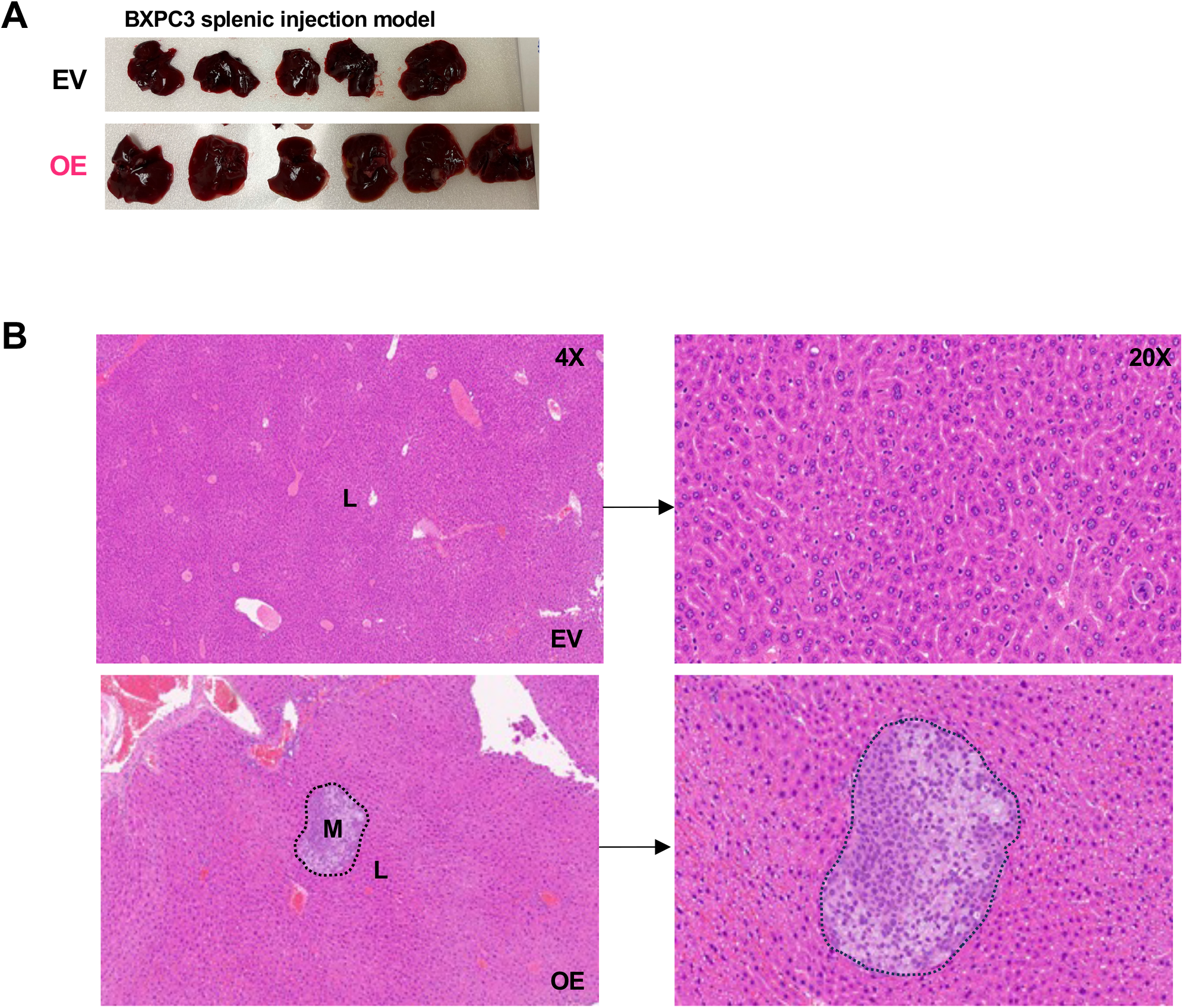
**A.** Representative images of liver metastatic burden after splenic injection of BxPc3 cells expressing EV or MICAL2-OE vectors into immunocompromised mice. **B.** Representative H&E images of livers shown in A. Normal liver tissue (L) and metastasis (M, dotted lines) are labeled.

**Supplementary Table 1.** Patient and tissue characteristics.

**Supplementary Table 2.** shRNA sequences used in this study.

**Supplementary Table 3.** siRNA sequences used in this study.

**Supplementary Table 4.** The sequences of oligonucleotides used for CDS cloning.

**Supplementary Table 5.** Primers used for q-PCR in this study.

**Supplementary Table 6.** The list of antibodies used for Western blotting.

**Supplementary Table 7.** SE-associated genes enrichment in tumor compared to normal tissues.

**Supplementary Table 8.** ASPC1 differential gene expression analysis in MICAL2-KD compared to scramble control cells.

